# From colonial clusters to colonial sheaths: analysis of *Microcystis* morphospecies in mesocosm by imaging flow cytometry

**DOI:** 10.1101/2023.07.07.545121

**Authors:** Adina Zhumakhanova, Yersultan Mirasbekov, Dmitry V. Malashenkov, Thomas A. Davidson, Eti Ester Levi, Erik Jeppesen, Natasha S. Barteneva

## Abstract

The alarming increase in the frequency of blooms of *Microcystis* in freshwater lakes and reservoirs occurs worldwide, with major implications for their ecosystem functioning and water quality. We applied FlowCAM-based imaging flow cytometry together with PCR and sequencing to get a comprehensive picture of the seasonal development of *Microcystis* community in a long-term running lake mesocosm experiment. The IFC analysis with manual taxonomic classification confirmed early findings with a machine learning algorithm that some *Microcystis* morphospecies completely disappeared and re-appeared along the mesocosm experiment timeline. This observation supports the hypothesis of the main transition pathways of colonial *Microcystis*. For the first time, colonial mucilaginous envelopes or sheaths were reported as separate entities, and not as a part of *Microcystis* colonies. The colonial sheaths may contain a few single Microcystis cells and reach significant numbers (thousands) during a cyanobacterial bloom. We also found that non-identifiable colonial small clusters of *Microcystis* cells are an important stage in the complex mosaic of a *Microcystis* bloom and are associated with the development of colonial forms. Our findings were validated by the principal component analysis coupled with the constructed associative matrices. We hypothesize that colonial sheaths may be crucial at *Microcystis* spp. dispersal and represent one of the stages of colonies development.

## 1. Introduction

Cyanobacteria are one of the most widely distributed and diverse phyla of bacteria, evolving around 2.5 billion years ago [1]. Globally, freshwater water bodies are increasingly threatened by cyanobacterial harmful algal blooms (HABs) due to anthropogenic pollution and climate change [2]. *Microcystis* spp. – potential toxic, bloom-forming cyanobacteria are one of the main cyanobacterial inhabitants of freshwater lakes and occur predominantly in a colonial form under natural conditions [3] but often as single or paired cells in a laboratory [4]. Colony formation mechanisms in *Microcystis* involve cell adhesion and cell division, where cells aggregate and divide, forming and encasing in an extracellular polymeric substance (EPS) layer. Colony formation of *Microcystis* with EPS provides a competitive benefit, including defense against grazing, density reduction, and dynamic streamlining [5]. Some morphospecies accumulate and release toxins and cyanopeptides in the surrounding water bodies, including microcystins (MCs) congeners [6]. Colonial morphology and size of *Microcystis* spp. are related to MCs production [7–9] and contribute to the success of this genus in freshwater ecosystems, leading to the global expansion and dominance in at least 108 countries and a majority of the continents [10–11]. Moreover, most *Microcystis* morphotypes could maintain their colonial characteristics for about one month in laboratory conditions [12]. Debates are ongoing on whether *Microcystis* morphospecies should be considered as individual species or as different morphoforms of single species and whether morphological diversity reflects phylogenetic diversity [12–13], and arguments depend on where species boundaries are set [14].

In shallow freshwater lakes, abiotic and biotic factors, including temperature, availability, and competition for nutrients, strongly affect phytoplankton succession and *Microcystis* seasonal blooms [6, 15–16]. *Microcystis* seasonal bloom goes through different phases and eventually leads to its decomposition and release of nutrients [17–18]. *Microcystis* bloom development has been considered in many mesocosms warming studies, often focusing on nutrient effects (N, N/P ratio) and microcystin synthesis and production [18–21]. The analysis of morphological characteristics of colonial *Microcystis* in published mesocosm and field studies is mainly done with light microscopy, though recently, the imaging flow cytometry (IFC) approaches with FlowCAM instrumentation have been used [22–24]. The IFC allows the high-throughput acquisition of thousands of colonies images in a relatively short period of time without substantial loss of information. Due to processing efficiency, the IFC imaging systems are well suited to analyze the heterogeneity of the *Microcystis* community and its functioning. Recently, we developed an IFC approach to differentiate the colonial morphology of five *Microcystis* morphospecies [25–26].

This study aimed to provide a detailed characterization of seasonal *Microcystis* development during a long-term running climate change mesocosm experiment. We hypothesize that different colonial *Microcystis* morphospecies represent at clusters related to the species’ development, multiplication stage, dispersal stage and originate from non-colonial small cellular clusters. The dispersal of *Microcystis* is supported by the formation of colonial mucilaginous envelopes-sheaths containing several single *Microcystis* cells and formed mainly from *M. wesenberghii*. The results are expected to contribute to understanding the *Microcystis* bloom development dynamics and provide new knowledge about how the dispersal of colonial *Microcystis* happens in nature.

## 2. METHODS

### 2.1 LMWE AQUACOSM mesocosm

The study was conducted as part of the Lake Mesocosm Warming Experiment (LMWE) at the facility owned by Aarhus University, located at Lemming, Central Jutland, Denmark (56°14’N, 9°31’E). The LMWE has been running continuously since 2003, and it is the longest freshwater lake mesocosm experiment in the world. The facility includes 24 outdoor freshwater cylindrical steel tanks of 1.9 m diameter and 1.5 m depth with ventilated paddles installed at the bottom, mixing the water to maintain uniform conditions. The water in mesocosms is sourced from groundwater with a water retention time of approximately 2.5 months [27]. The temperature regimes include unheated control with ambient water temperature (AMB) and two elevated temperature settings based on the Intergovernmental Panel on Climate Change (IPCC) climate scenarios for the period 2071-2100, IPCC A2 (*ca.* +3 °C) and IPCC A2+50% (*ca.* +4.5 °C), with four replications for each regime. The water temperature is maintained in the heated tanks using an automatic heating system with three electrically powered steel heating elements (750 W each). Unheated mesocosms were equipped with imitation of heating elements. Tanks D1, F1, G1 reflect the ambient water temperature, while the next two subgroups represent two IPCC scenarios regarding water temperature in the lake by the end of the century (A2 and A2 + 50%). Tanks D2, F2, G2 imitate a water temperature setting that is 2 - 3 ^0^C warmer than seasonal water temperatures in the lakes (A2), and tanks D3, F3, G3 demonstrate the model of water temperature setting that is 5 - 6 ^0^C higher than the average seasonal water temperature in the lake (A2 + 50%) (**Figure 1**). A2+50% tanks were excluded from further consideration due to the negligible quantity or absence of *Microcystis* colonies. The current study was run in 6 mesocosms with a high nutrient level which has been constantly supplied with 108.6 mg N per m3 per day and 2.7 mg P per m3 per day in the form of Ca(NO_3_)_2_ and Na_2_HPO_4_, respectively. Each mesocosm was artificially supplied with 2152 mg N and 54 mg P per week in addition to the nutrient inputs from groundwater of approximately 2-5 mg P and 30-63 mg N per week.

**Figure 1.**
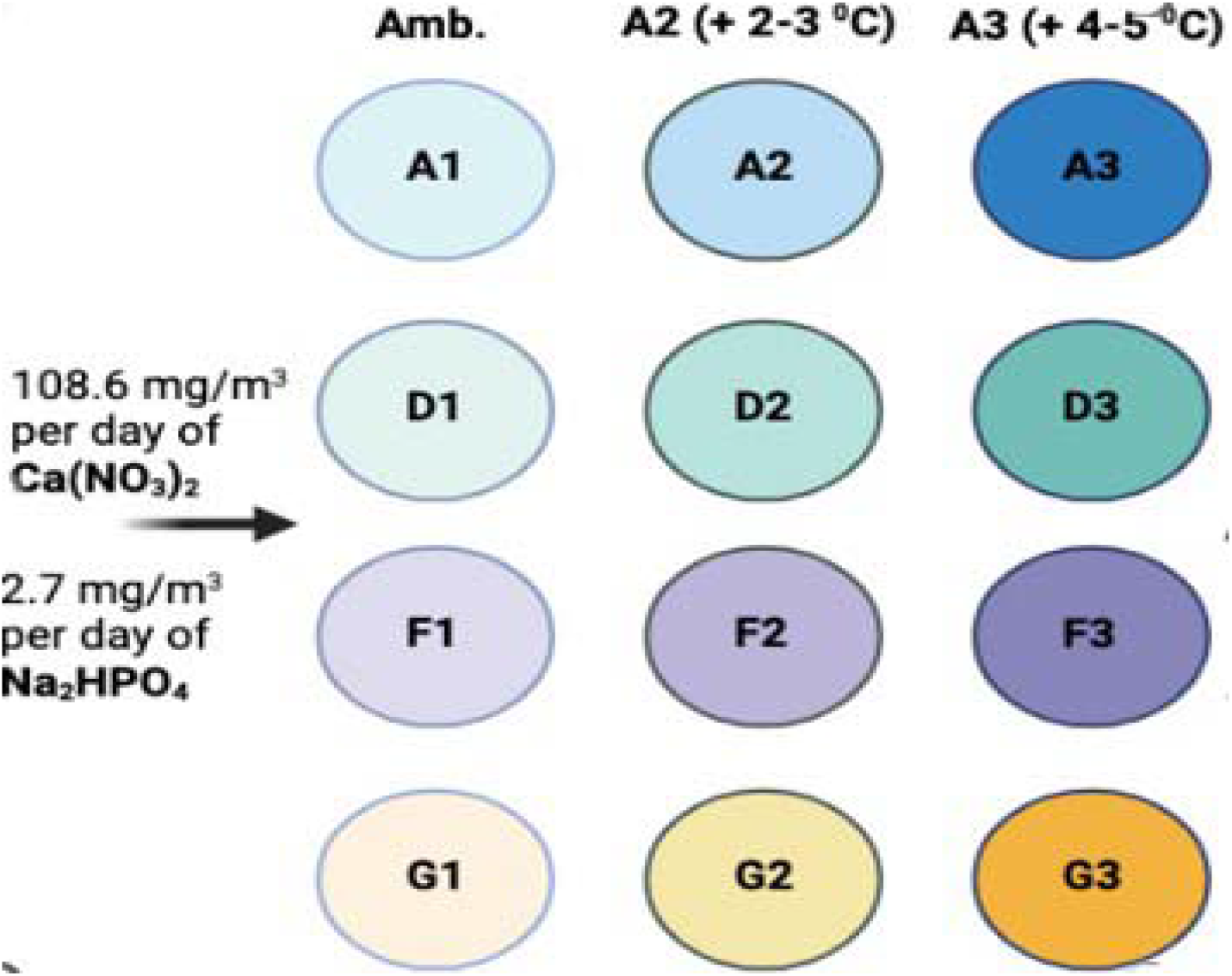
Schematic representation of mesocosm tanks used in this study.

### 2.2 Samples acquisition

Integrated water samples were collected using a 1 m long tube water sampler on 13 different dates from the 23^rd^ of May to the 17^th^ of September 2019, resulting in 117 samples. All samples were fixed with 1 % glutaraldehyde. The following parameters were measured and used as independent variables: water temperature (^0^C), turbidity, total phosphorous (P_total_, mg/L), inorganic phosphorous (PO_4_, mg/L), total nitrogen (N_total_, mg/L), nitrate + nitrite (NO_2+3_N, mg/L), ammonium (NH_4_, mg/L). To analyze the seasonal succession of colonial *Microcystis* spp. in 2019 summer, tanks D, F, and G were used with their corresponding sub-units.

### 2.3 FlowCAM-based Imaging Flow Cytometry

The imaging analysis was performed using the FlowCAM VS-4 (Yokogawa Fluid Imaging Technologies, USA), an imaging flow cytometer that combines the properties of a microscope and flow cytometer. 10 mL from each undiluted sample was filtered with a 100 µm mesh and recorded with 10x and 20x objectives and a 100 µm flow cell. Particles were captured in autoimage mode with 20 frames per second rate, including the collection of binary images. The flow rate was set at 0.3 mL/min, which resulted in 33 minutes of recording of the field of view for each sample. In this study, the autoimage operating mode in FlowCAM (Yokogawa Fluid Imaging Technologies, USA) was utilized as it has an optimal resolution size limit for research focused on *Microcystis* spp. [28] Light and dark pixel thresholds were set at 20 and 40, respectively, to intensify the particles over the background. The VisualSpreadsheet vs. 4.0 software (Yokogawa Fluid Imaging Technologies, USA) was used to was used to classify taxonomic classes of interest and abundances manually.

### 2.4 Morphological classification of Microcystis spp. morphospecies and sheaths

Five major *Microcystis* morphospecies were targeted for classification: *M. novacekii*, *M. ichtyoblabe*, *M. smithii*, *M. aeruginosa*, *M. wesenbergii* [25–26, 29] (**Figure 2**).

**Figure 2.**
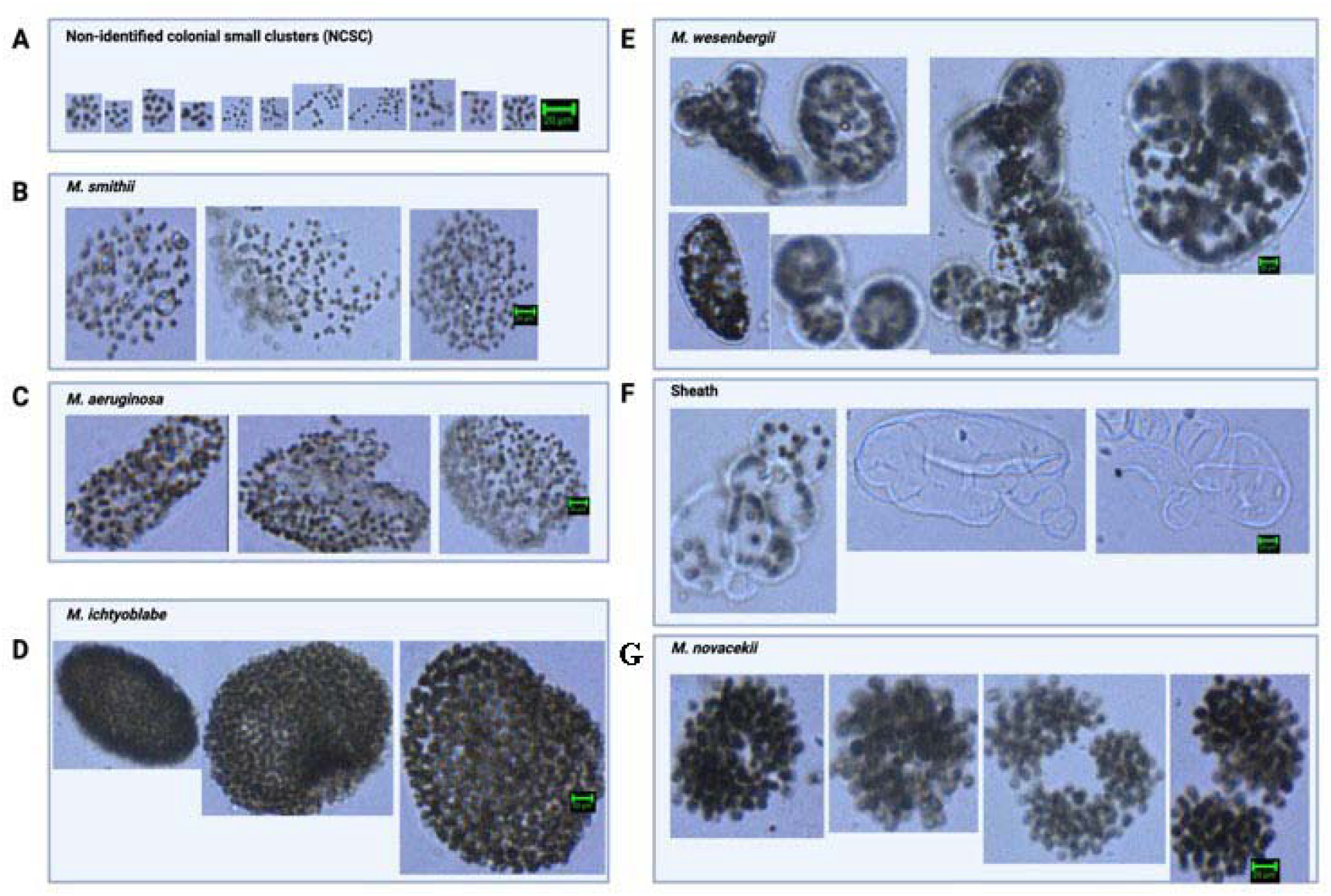
Gallery of *Microcystis* spp. Images were identified and used for morphological classification via the FlowCAM imaging system. A. Non-identified colonial small clusters (NCSC); B. *M. smithii*; C. *M. aeruginosa*; D. *M. ichtyoblabe*; E. *M. wesenbergii*; F. colonial sheaths mirroring *M. wesenbergii* shape; G. *M. novacekii*.

Additionally, colonial mucilaginous envelopes with many, a few, or none *Microcystis* cells were classified separately and marked as sheaths. Moreover, on some dates, we detected non-identified colonial small clusters of *Microcystis* spp. cells (NCSC). The taxonomic classification was performed manually with VisualSpreadsheet vs.4.0 software, and relative abundances (particles/mL) of *Microcystis* morphospecies and sheaths were used in the data analyses. Sheaths were further divided into three sub-categories: type A, type B, and type C, according to the quantity of *Microcystis* cells inside the sheath. Types A, B, and C contain sheaths with a number of cells from 0-10, 10-50, and >50, respectively. The examples of sheath sub-categories and colonies of *M. wesenbergii* are demonstrated in **Figure 3**.

**Figure 3.**
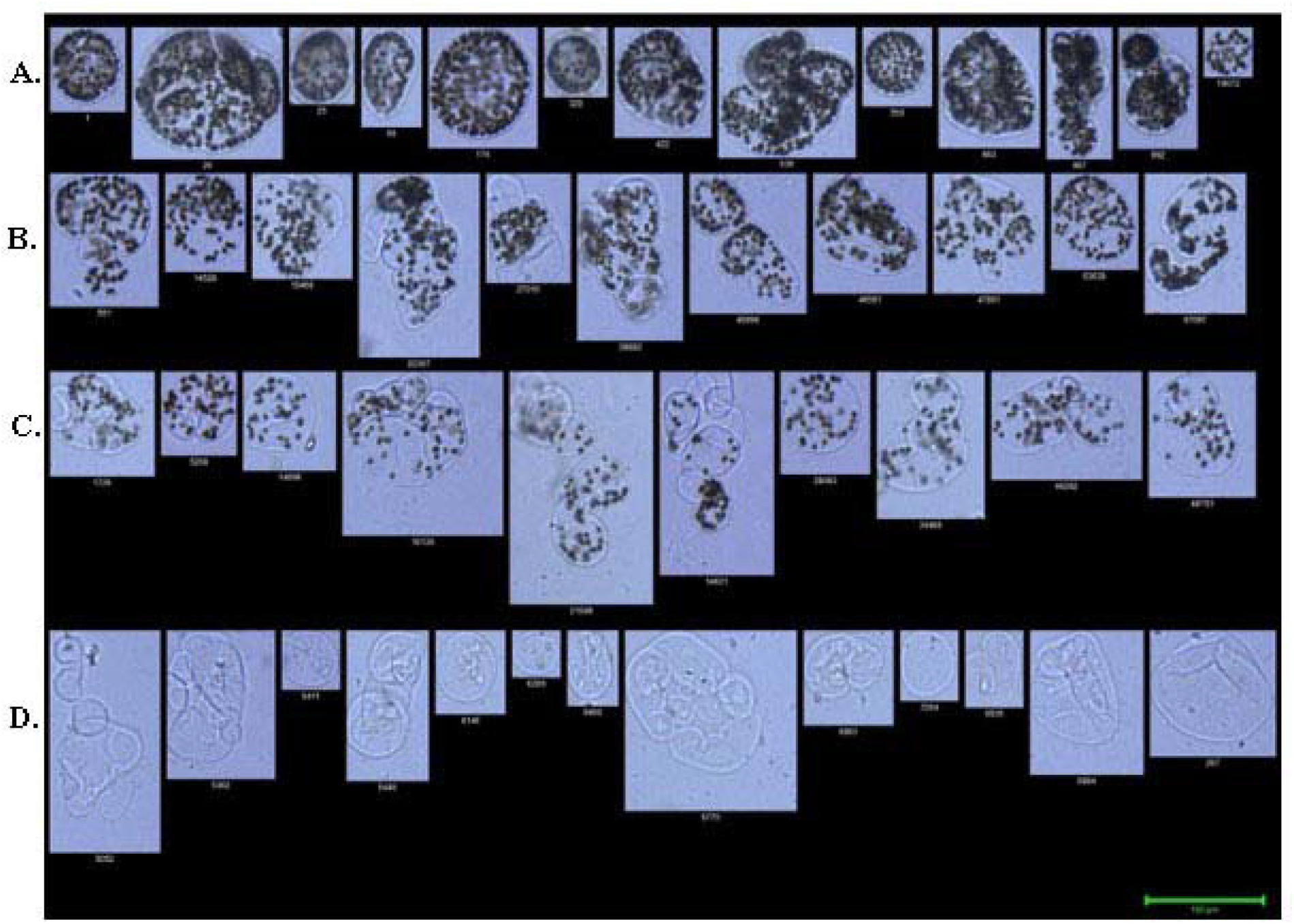
Classification of *M. wesenbergii* colonies and colonial sheaths sub-categories from LMWE mesocosm 2019. **A**. *M. wesenbergii*; **B**. type C sheaths (more than 50 cells per colony); **C**. type B sheaths (10-50 cells); **D**. type A sheaths (less than 10 cells, including empty sheaths).

### 2.5 Light microscopy

Taxonomic identification of *Microcystis* colonies was confirmed by an experienced taxonomist (co-author Dmitry V. Malashenkov) based on distinctive morphological features of colonies and cells [29–30] using a Leica DM2500 microscope (Leica Microsystems, Germany) equipped with differential interference contrast (DIC) and 10x, 20x, 40x, 63x, and 100x objectives. Randomly selected colonies were measured at 100x, 200x, and 400x magnifications.

### 2.6 PCR analysis for Microcystis spp. and microcystin production gene

For molecular biological analysis, 1 liter of water sample was collected from 8 different mesocosm tanks. Samples were entrapped in the filter membrane with a pore size of 0.22 μm and stored in conical Falcon tubes (BD Biosciences, USA) at 4°C. The DNA was extracted by the manufacturer’s recommendations using the PowerWater DNA Isolation Kit (Qiagen, USA). The DNA extract with a final volume of 100 μL was stored at -20°C. The PCR reactions were performed to target the *Microcystic*-specific 16S rRNA gene for phylogenetic identification of *Microcystic* spp. Also, the *Microcystis*-specific *mcyE* gene was targeted to identify potential toxin production. According to Pacheco and co-authors [31], the *mcyE* gene had the highest positive correlation with actual microcystin concentration among other genes within the microcystin biosynthesis gene cluster (*mcyA*, *mcyB,* and *mcyD*). The primer design for the *Microcystis*-specific *mcyE* gene was based on the work of Rudi and co-authors, and Vaitomaa and co-authors, respectively [32–33]. The information on the primers used is summarized in **Table 1** below. These primers were synthesized by Evrogen company (RF).

**Table 1.**
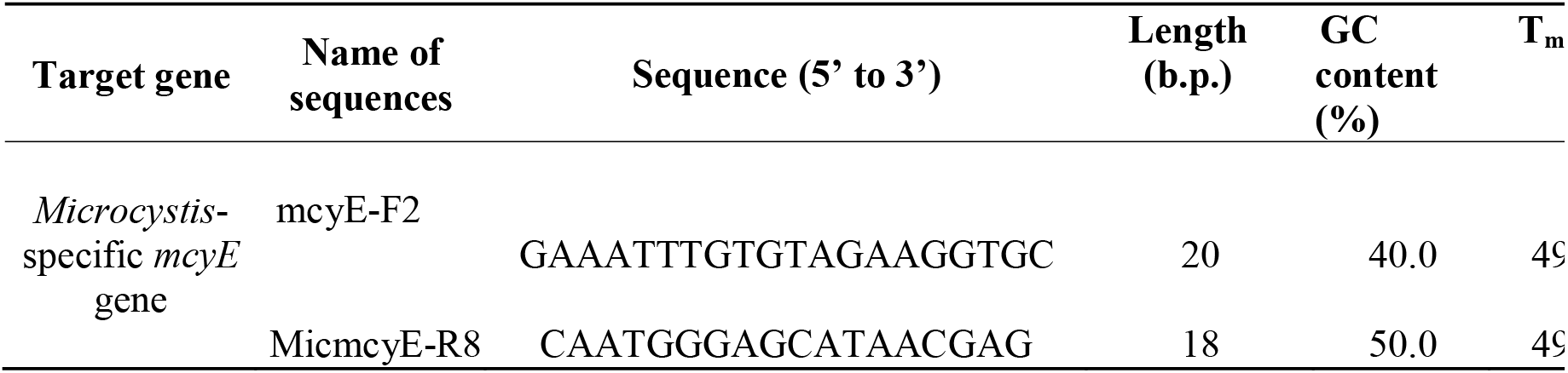
List of primers used in the study.

### 2.7 PCR settings

The PCR reaction mixture contained DNA sample (about 3 μL), DreamTaq Hot Start PCR Master Mix (25 μL), 10 μM forward primer (0.5 μL), 10 μM reverse primer (0.5 μL), nuclease-free water, and these reaction tubes were placed on a thermocycler (Bio-Rad, USA). Reactions started with the initial denaturation step (3 min, 95°C), which was followed by 40 cycles of denaturation (30 sec, 95°C), annealing (30 sec, depending on T_M_ of primers), and extension (60 sec, 72°C) steps. The final extension was set for 5 min at 72°C. Products of PCR reactions were stored at -20°C freezer. These products were visualized by SYBR^TM^ Safe DNA stain (Life Sciences, USA) in 1% agarose gel electrophoresis.

### 2.8 Sanger sequencing

Selected samples were sequenced by Evrogen company (RF), and the nucleotide composition of obtained sequences was used for a search in the NCBI Nucleotide collection database (USA).

### 2.9 Statistical analysis

Data were checked for normality via D’Agostino & Pearson and Shapiro-Wilk tests. Then Principal Component Analysis (PCA) biplots and Spearman correlation matrixes were calculated to identify significant correlations and associations between environmental parameters and *Microcystis* morphospecies pairs using GraphPad Prism vs. 9.0 software (Dotmatics, USA). The difference in community composition (date-to-date comparison) was checked via ANOSIM test. The Kruska-Wallis and Bonferroni-Dunn’s tests were used to identify statistically significant changes in the diameters of colonial sheaths (p-value < 0.05).

## 3. Results

### 3.1. Seasonal variations in Microcystis spp. abundance

Distribution and spatial-temporal seasonal variations of *Microcystis* morphospecies from May to September 2019 are shown in **Figure 4**. The recurring patterns of seasonal succession of *Microcystis* spp. were demonstrated, i.e., morphospecies could be totally absent on one date and present in a great number on the following date or even after several weeks. Generally, tanks with A2 water temperature scenarios had a higher diversity of *Microcystis* spp. and two or more bloom events. Numerous *Microcystis* bloom events consisted of *M. smithii*, *M. aeruginosa, M. ichtyoblabe. M. wesenbergii*, and *M. novacekii* colonial morphoforms, as well as NCSC and sheaths. Colonies of *M. ichtyoblabe. M. aeruginosa,* and *M. smithii* were never found to be dominating morphospecies in any of the detected blooms.

**Figure 4.**
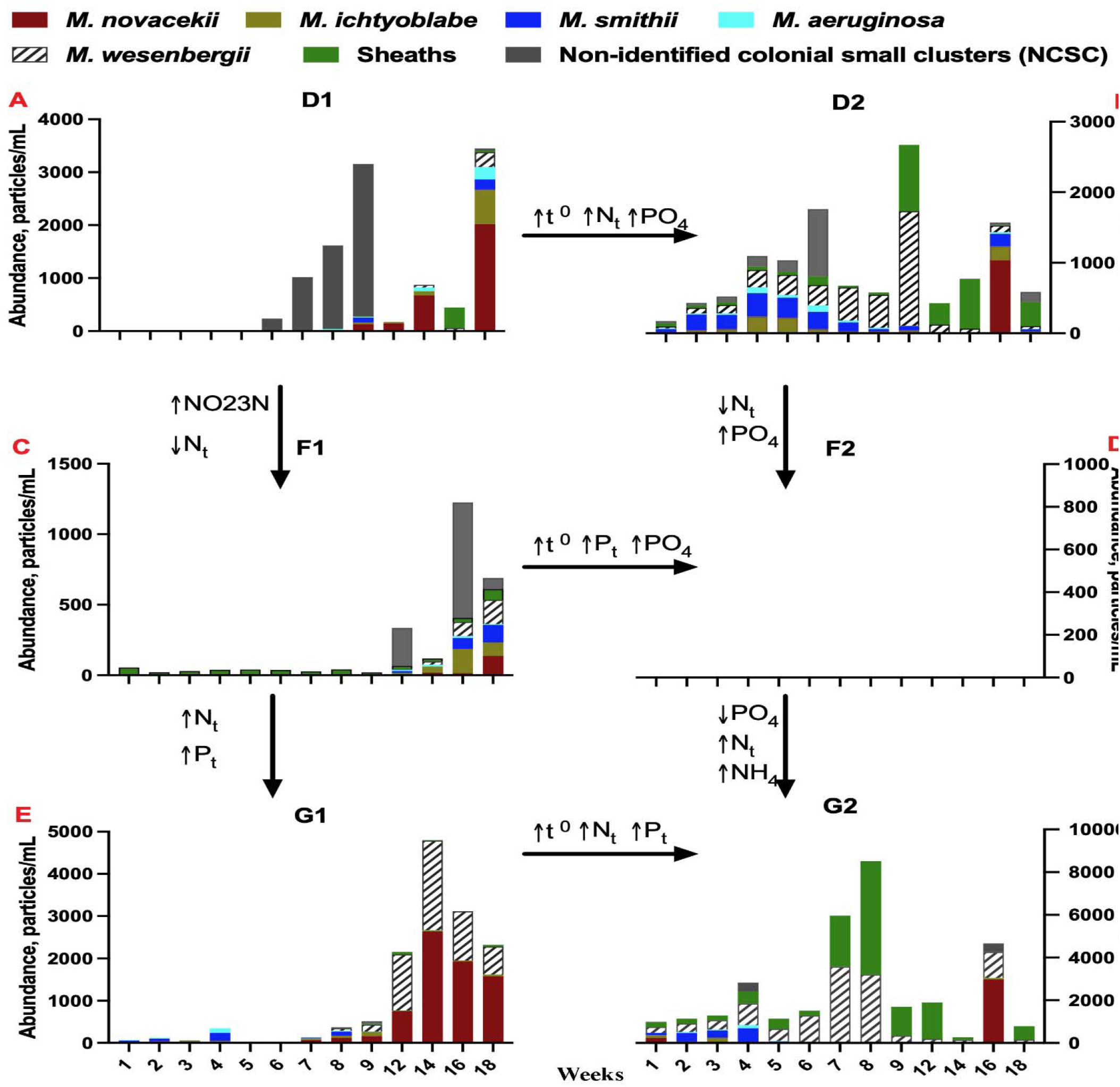
The abundance of M*icrocystis* morphospecies in the LMWE-2019 experiment in tanks **A.** D1; **B.** D2; **C.** F1; **D.** F2; **E.** G1; **F.** G2 (particles/ml states for colonies/ml). No *Microcystis* sp. colonies in F2 tank but centric diatoms (*Stephanodiscu*s sp.; May-July) and *Micractinium pusillum* (July to August 2019) were found.

### 3.2. Sequencing analysis of PCR products

The *Microcystis mcyE* gene was found in samples from most of the analyzed mesocosm tanks (**Suppl. Tables 1** and **2**). These results concur with data from FlowCAM analysis of different *Microcystis* morphospecies (**Figure 4**).

### 3.3. Influence of temperature and nutrients on Microcystis spp. abundance

Water temperature variations were similar for tanks that followed one or another water temperature simulation: ambient or A2. For tanks D1, F1, G1, blooms were detected at or after the date when the water temperature peaked (1st of July = ∼21^0^C). Higher temperature settings (A2 tanks) correlated with a greater diversity of detected morphospecies from the beginning of the monitored period as well as with earlier observations of bloom events. We observed a positive correlation between the date-to-date shifts in water temperature and the abundance of *Microcystis aeruginosa* colonies (**Figure 5**). The overall increase in water temperature might be the driving force for the sequential development of colonial *M. wesenbergii* morphospecies and sheaths rather than changes in water temperature between dates (**Figure 5**). Our results suggest that the initial threshold of water temperature for growth was around 21^0^C after which *Microcystis* spp. initiated a series of bloom-forming events.

**Figure 5.**
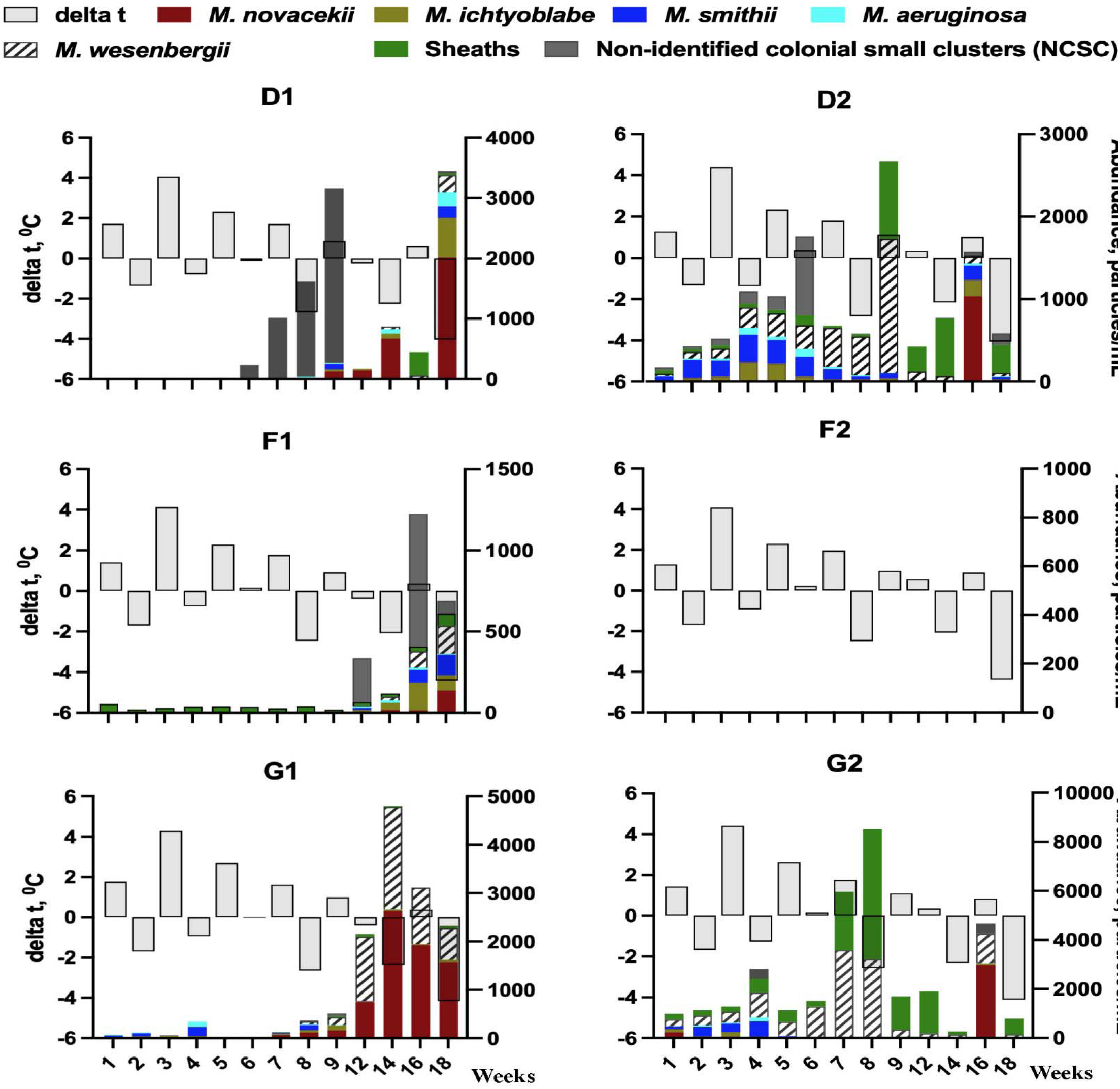
Seasonal variations of the water temperature (Δt – shift in water temperature,^0^C) and abundance of colonial and non-colonial *Microcystis* spp. morphospecies from May to September 2019 in tanks D1, D2, F1, F2, G1, G2.

The variations of measured concentrations of nutrients are presented in **Suppl. Figures 1 and 2**. Colonies of *M. novacekii* were found even if N_total_ had potential limiting concentrations (3rd of September for all tanks). Also, *M. novacekii* colonies were present at concentrations of P_total_ of at least 0.5 mg/L (tanks D1, G1, D2, G2). Colonies of *M.aeruginosa* emerged in early summer periods or mid-summer at decreasing water temperature.

### 3.4. Seasonal variations in abundance of M. wesenbergii and sheaths

Type A sheaths, and *M. wesenbergii* were detected independently; however, type B and C were found when either type A sheaths or *M. wesenbergii* colonies were present. The variety of sheaths was observed with an increase in water temperature (D2 vs. D1, G2 vs. G1). Generally, a higher water temperature setting favored the presence of type A sheaths (tanks D2 and G2) **(Figure 6)**.

**Figure 6.**
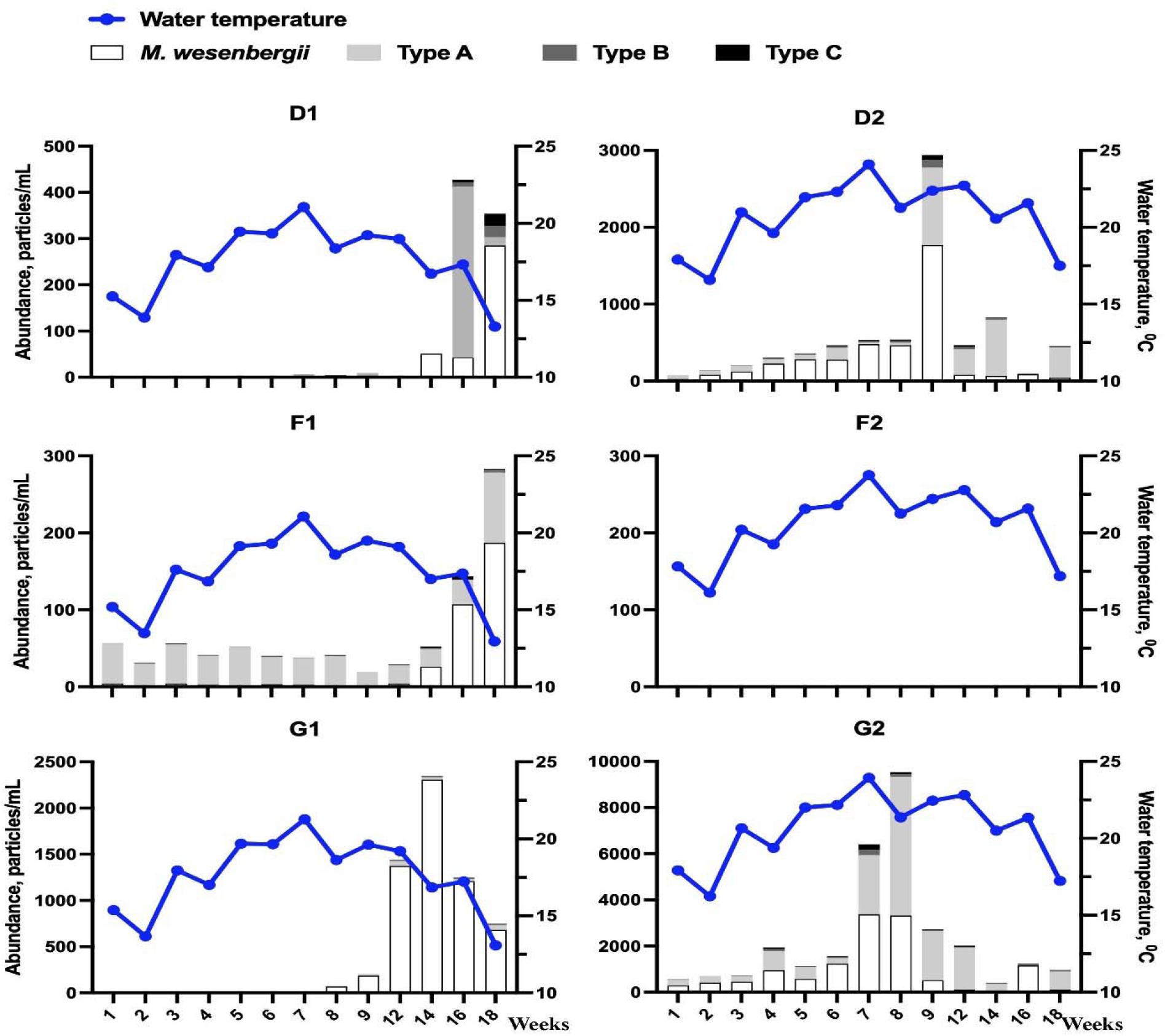
Seasonal variations of *M. wesenbergii* and sheath types (A, B, C) versus water temperature from May to September 2019 in tanks D1, D2, F1, F2, G1, G2.

The growth of the number of *M. wesenbergii* colonies coincided with an increase in the concentrations of N_total_ and P_total_ in ambient temperature tanks (**Figure 7**). However, in tanks D2 and G2, the maximum magnitude of *M. wesenbergii* colonies and sheath types was in the middle of the summer (1st of July), matching the peak of water temperature, indicating that increased water temperature may be a principal factor over the concentration of N_total_ and P_total_ as soon as it reaches some optimum threshold. From the data, the minimum concentration of P_total_ at which bloom consisting of *M. wesenbergii* was observed was 0.36 mg/L.

**Figure 7.**
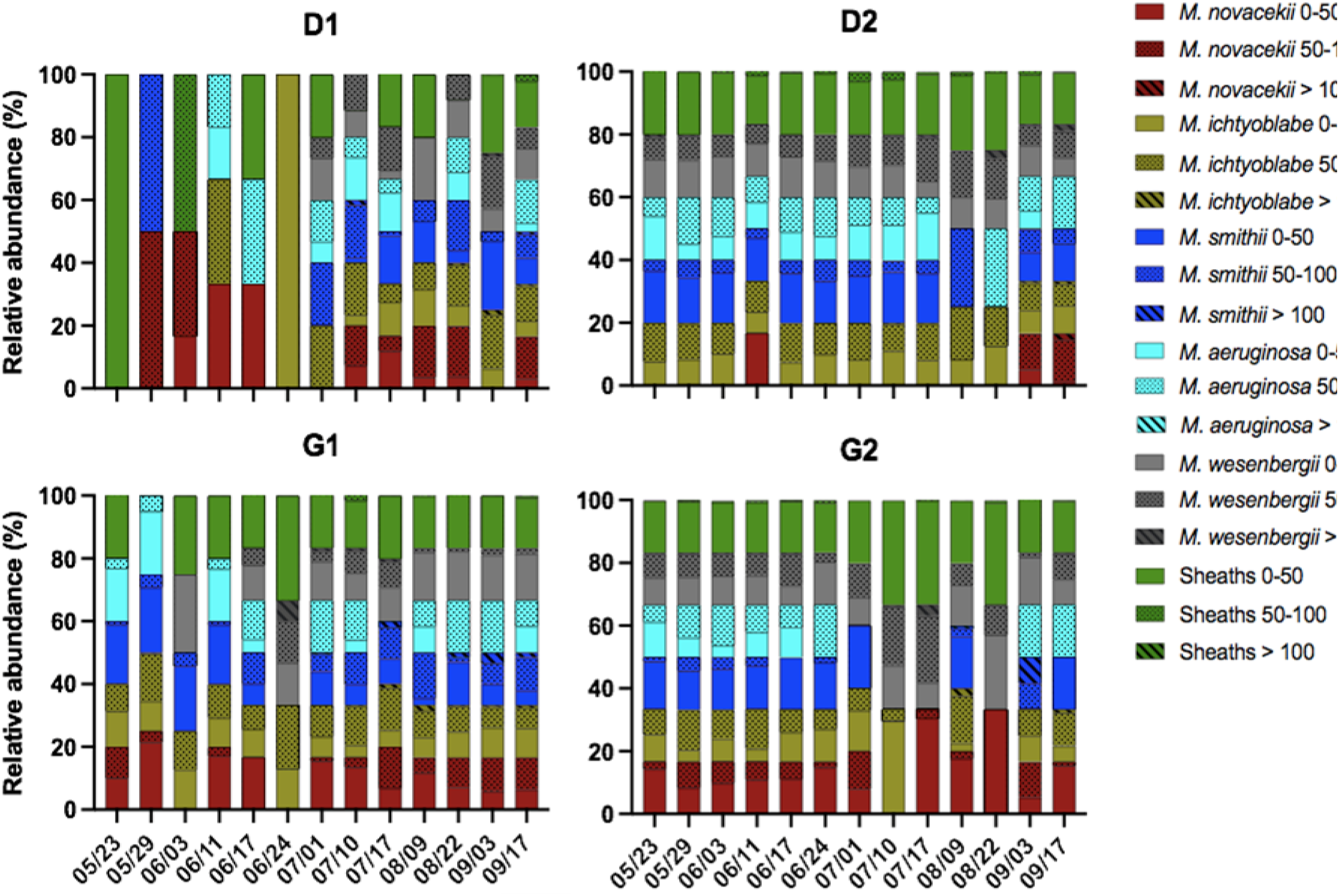
The relative contribution of *Microcystis* spp. morphospecies to colonial *Microcystis* community and sheaths (total is 100 %) in tanks D1, D2, G1, G2. Colonies are divided into three subtypes according to ABD parameter (0-50, 50-100, > 100). Groups with ABD between 0 and 50 are represented with rigid colors, groups with ABD 50-100 have dotted colors, and groups with a diameter of more than 100 µm have diagonal lines.

Multiple and colonies-rich blooms appearance of predominantly *M.wesenbergii* colonies and sheaths usually happened in the mid- and late-summer period when the water temperature was at the highest level. However, under the right conditions (e.g., an increase in temperature), colonies of *M. wesenbergii* and sheaths formed in early summer or even in spring (tanks D2 and G2). Accompanying sheaths were often found to contain various numbers of *Microcystis* cells coupled with high polysaccharide content (**Figure 2**). Sheaths of type B and type C but not type A were co-present with *M. wesenbergii* colonies or with type A.

### 3.5. The changes in diameter (ABD) of colonial morphospecies and sheaths

Area-based diameter (ABD) was used to allocate colonial *Microcystis* morphospecies and sheaths into three groups (**Figure 7**). In most cases, a higher temperature setting (D2 versus D1, G2 versus G1) seemed to have a positive effect on the balance of all three size (ABD parameter) groups. Tanks D2 and G2 had fewer fluctuations of ABD parameter for all colonial morphophorms and sheaths in contrast to D1 and G1.

The dynamic of colonial sheaths is shown in **Figure 8**. Besides the stronger magnitude of blooms, we observed a higher number of blooms-forming events at higher temperature (comparing tanks D2 vs. D1 and G2 vs. G1).

**Figure 8.**
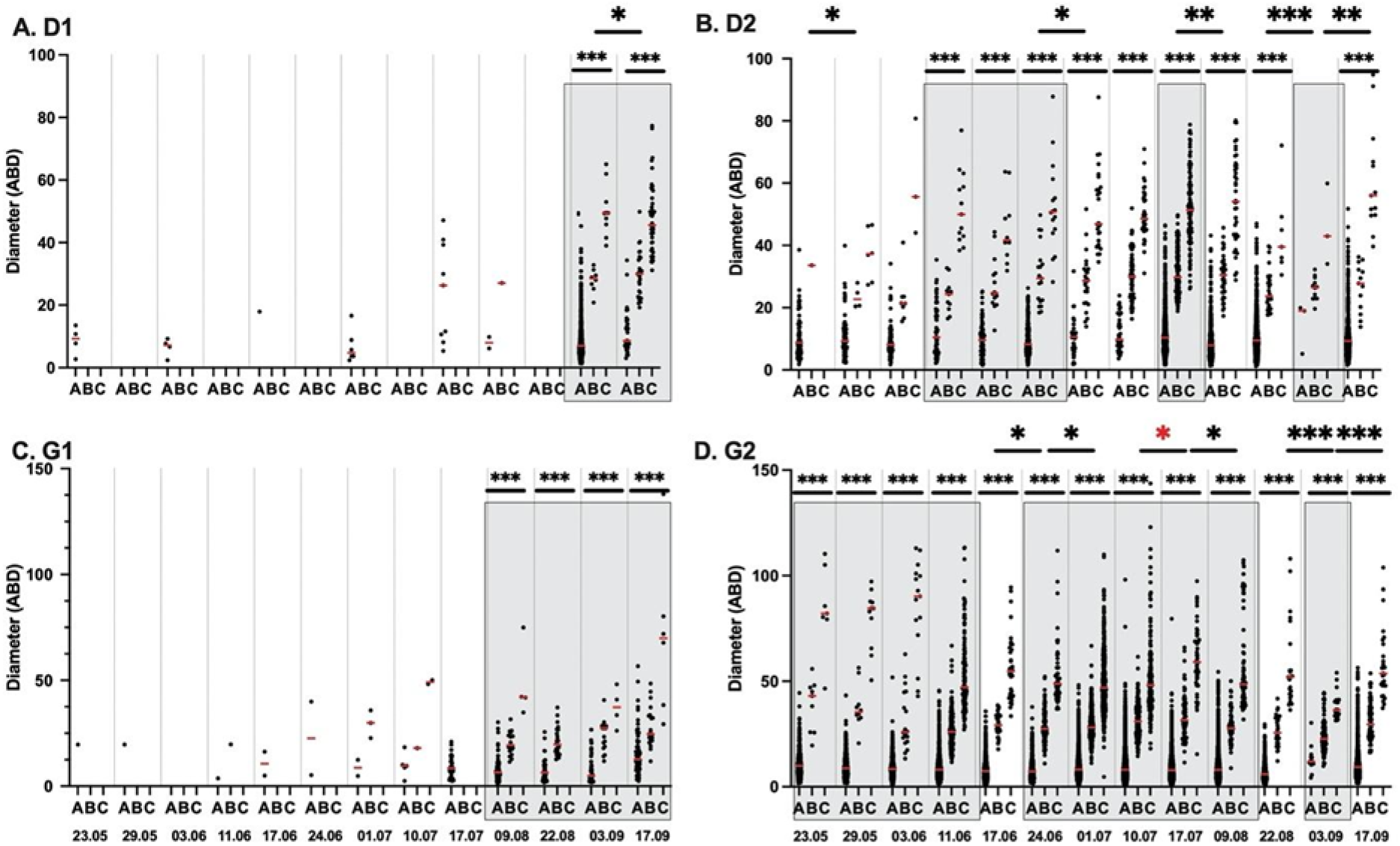
The dynamic of sheaths size (ABD) in tanks D1, D2, G1, G2. The lower asterisks represent differences between either type of sheaths or between diameters of sheaths and *M. wesenbergii* colonies (p<0.05). The upper asterisks represent differences in community composition (date-to-date comparison) checked via ANOSIM test (p<0.05). **A.** D1 tank; **B.** D2 tank; **C.** G1 tank; **D.** G2 tank.

### 3.6. Seasonal succession of Microcystis spp. community: PCA and correlation matrices

The multivariate associations are presented in the form of PCA biplots and Spearman correlation matrices for the tanks where a substantial amount of *Microcystis* spp. was found (D1, D2, F1 in **Figure 9A**, and G1, G2 in **9B**). The dataset includes various physico-chemical variables and classified *Microcystis* morphotypes.

**Figure 9.**
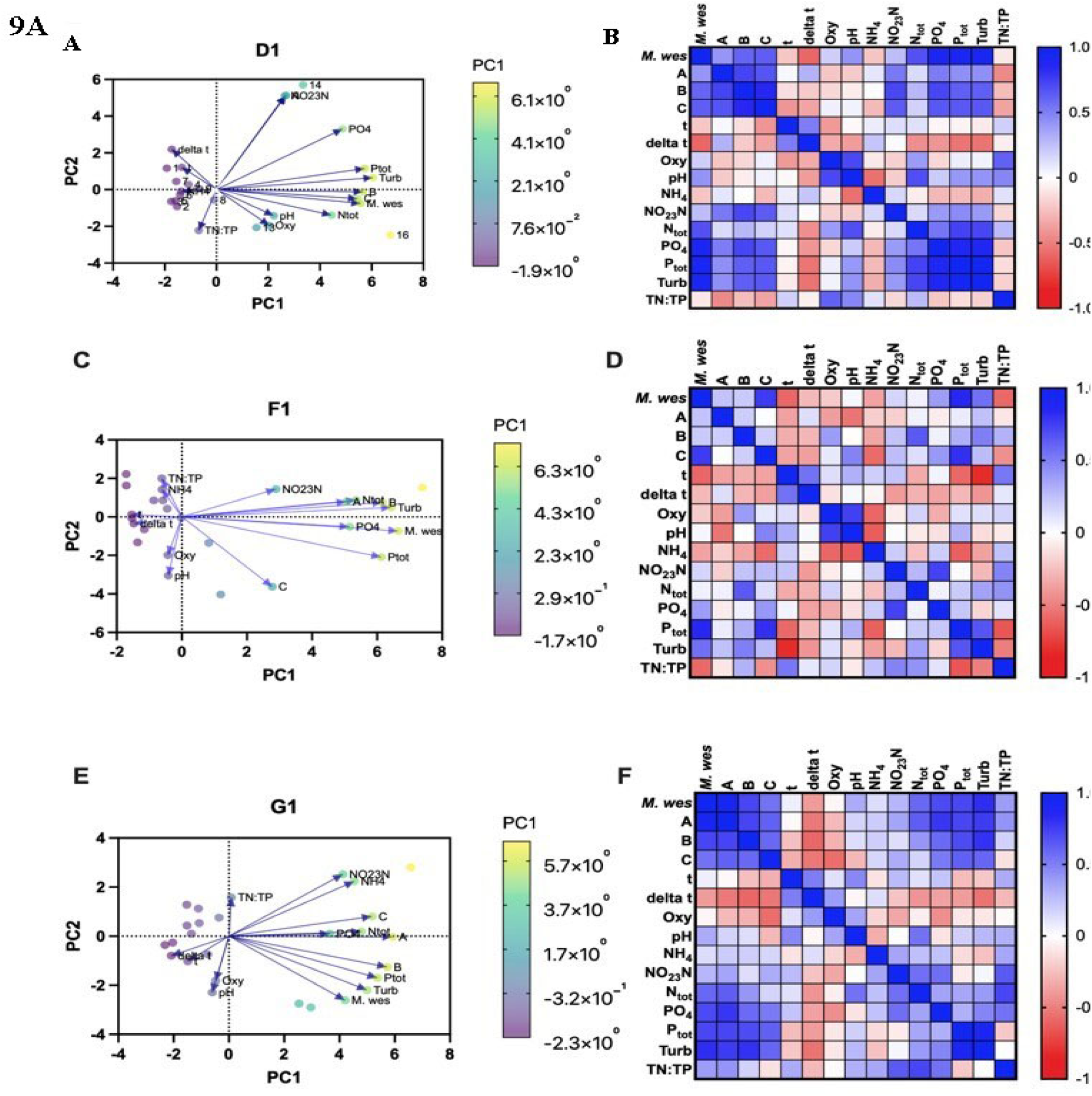
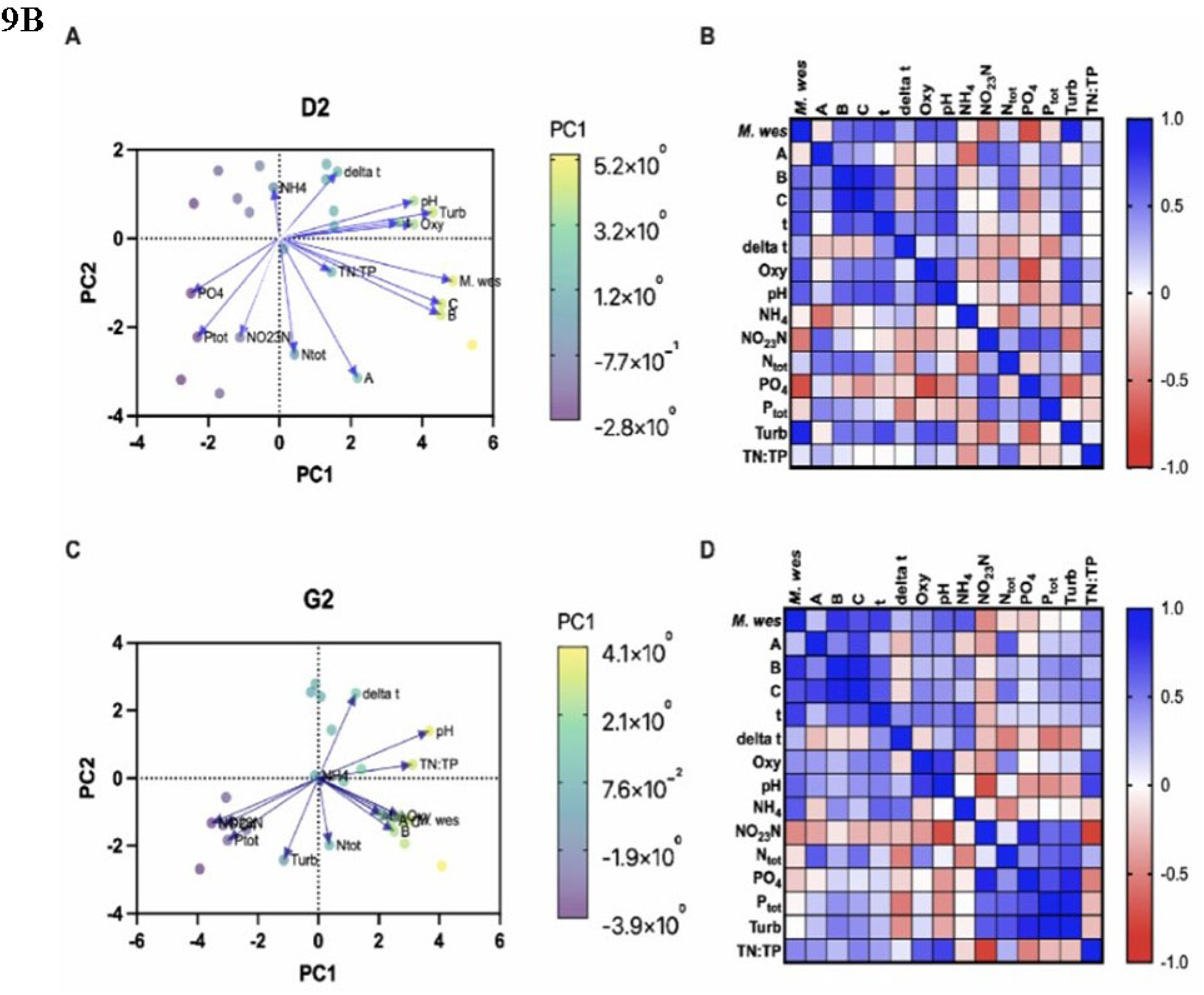
PCA biplot of normalized data and Spearman correlation matrices for *M. wesenbergii* and type A, B, C sheaths and environmental parameters for **A.** tanks with ambient temperature D1, F1, G1; and **B.** tanks D2 and G2. The blue color indicates a strong positive correlation, and the red color indicates a strong negative correlation. Significant correlations (p-value <□0.05) are labeled by “*.”

It is important to note that PCA and constructed matrices do not indicate linear relationships but rather associations between analyzed groups. Considering the limitations of the PCA analysis, we are interpreting **Figures 9A,B** as follows. Firstly, changes in water temperature settings caused changes in the relationship between morphospecies. Furthermore, higher nutrient loading strengthened associative links between morphospecies. At warmer water temperatures, the correlation relationships between *Microcystis* morphospecies, except *M. wesenbergii* and sheaths, became reversely associated. These reverse associations might lead to the competitive advantage of *M. wesenbergii* and sheaths compared to other morphospecies. Spearman correlation did not show any significant associations between the form of N loading and the growth of *Microcystis* spp. colonies (**Figures 9A,B**).

PCA analysis of trophic preference showed no uniform correlation between phosphorous (P_total_, PO_4_) and the abundance of various morphospecies. In tanks with lower temperature regimes, the correlation was mainly positive. Finally, we also found that the concentration of phosphorus (P_total_) appears to be an important factor for NCSC that decreased their quantity, possibly, by forcing to transform into larger, identifiable colonies.

## 4. Discussion

*Microcystis* morphospecies repeatedly occurs in the seasonal succession from spring to fall in certain lakes and areas [16, 34–36]. The primary focus of the published studies on *Microcystis* has been mainly on the *Microcystis aeruginosa* complex [37–38]. Recently, interest shifted to other *Microcystis* morphospecies, causing debates about whether they are distinct species or closely related monophyletic clades [39–40].

Here we studied long-term running mesocosm mimicking the functionality of a natural ecosystem to elucidate the seasonal transitioning of *Microcystis* morphospecies. Previous works from our laboratory confirmed the effectiveness of FlowCAM (Yokogawa Fluid Imaging Technologies, USA) IFC as a tool for analyzing mesocosm samples for a substantial period, e.g., three-four months of bi-weekly monitoring routine [25–26, 41]. We used FlowCAM autoimage operating mode, which has an optimal resolution size limit for research focused on *Microcystis* spp. [28]. Applying FlowCAM-based IFC combined with PCR and sequencing, we could identify the presence of *Microcystis* sp. even in samples with a low concentration of the cyanobacteria.

### 4.1 Temperature effect on Microcystis colony size and abundance

There is a common agreement that elevated temperatures induce larger colonies, increase toxicity, and contribute to the development larger blooms of *Microcystis* spp. [42–44]. Our study indicates that higher temperature settings cause a greater intraspecific variety of morphospecies (D2/D1 and G2/G1 tanks). We found that the optimal water temperature for the growth of *Microcystis* colonies ranged from 20^0^C to 25^0^C, which is in agreement with the optimum 25-27^0^C observed in previous studies of *Microcystis* spp. [45–46]. Increasing water temperature not only led to higher diversity in *Microcystis* spp. population, but also favored more frequent bloom-forming events in tanks D2 and G2. Regarding *M. aeruginosa* colonies, several research groups [47–48] demonstrated that they increase in abundance with a decline in water temperature, which concurs with our observations. Also, we found the co-occurrence and association between *M. ichtyoblabe*, *M. aeruginosa*, and *M. smithii* colonies was stronger in the high-temperature tanks independent of nutrient levels. In contrast, nutrients apparently were a more prominent factor for the enumeration of these colonies in tanks with lower temperature settings.

### 4.2 Seasonal Microcystis morphospecies succession

Our results revealed the periodic nature of *Microcystis* colonies’ development throughout May - September of 2019: some of the morphospecies can be undetectable at one date and be in a significant number or even participate in the formation of bloom at the other date. One possible reason explaining such rapid variations in the abundance might be the distinctive appearance of the intermediate morphospecies with a high level of phenotypical plasticity [25, 49].

NCSC refers to small aggregations of *Microcystis* cells and is not a transitional form but a crucial stage in developing the *Microcystis* community [50, 51]. Early Wu and Kong [52] pointed out the uniform distribution of small colonies in waters under turbulent mixing by wind gusts. Still, the highest abundance of small colonies is to be found in the middle layers under extensive light radiation. Smaller colonies or small aggregations of *Microcystis* cells are often discovered in a colder period of the year [53], which correlates with our findings. According to the meta-analysis made by Xiao and co-authors [54], it is likely that *Microcystis* small grouped aggregated cells start with adhesion to each other leading to the formation of *M. ichtyoblabe*-like, *M. aeruginosa*-like, and *M. smithii*-like colonies. One of the intriguing results of our study is the strong association between *M. ichtyoblabe*-like, *M. aeruginosa-*like, and *M. smithii*-like colonies. These morphotypes’ general co-occurrence suggests they belong to the same colonial cluster. Another possible explanation is the temporal appearance of these morphotypes, mainly after NCSC blooms or during them.

Our findings raise a question regarding the transition of *Microcystis* spp. cells to a secondary colonial structure. We suggest that the observed five *Microcystis* morphospecies are part of single species with remarkable morphological plasticity. Early Šejnohová & Maršálek [55] pointed out a possible clustering within *Microcystis* species based on morphological and molecular markers. Moreover, Xiao and co-authors [54] proposed that the colony formation of *Microcystis* spp. cells start with the same morphotype and then change in response to environmental stressors.

Based on the observed dynamics of *Microcystis* morphospecies, NCSC, and colonial sheaths in this mesocosm experiment, we hypothesize that under limiting nutrient conditions and at ambient water temperatures, overwintered *Microcystis* cells either form pre-colonial structures and then proceed to bloom-forming events or exist as small colonial clusters after overwintering in the surface sediment (**Figure 10**). As a part of our hypothesis, we propose that *M. ichtyoblabe*-like and *M. smithii*-like colonies develop from NSCS. However, the nature of relationships between NCSC – *M. ichtyoblabe*, NSCS – *M. smithii* might be ambiguous. It appears that *M. aeruginosa*, *M. ichtyoblabe*, and *M. smithii* belong to one initial cluster while sheath-forming *M. wesenbergii* is more specialized in the proliferation and spreading of cells/colonies, and *M. novacekii* with NCSC are necessary for the renewal of this *Microcystis* spp. cycle.

**Figure 10.**
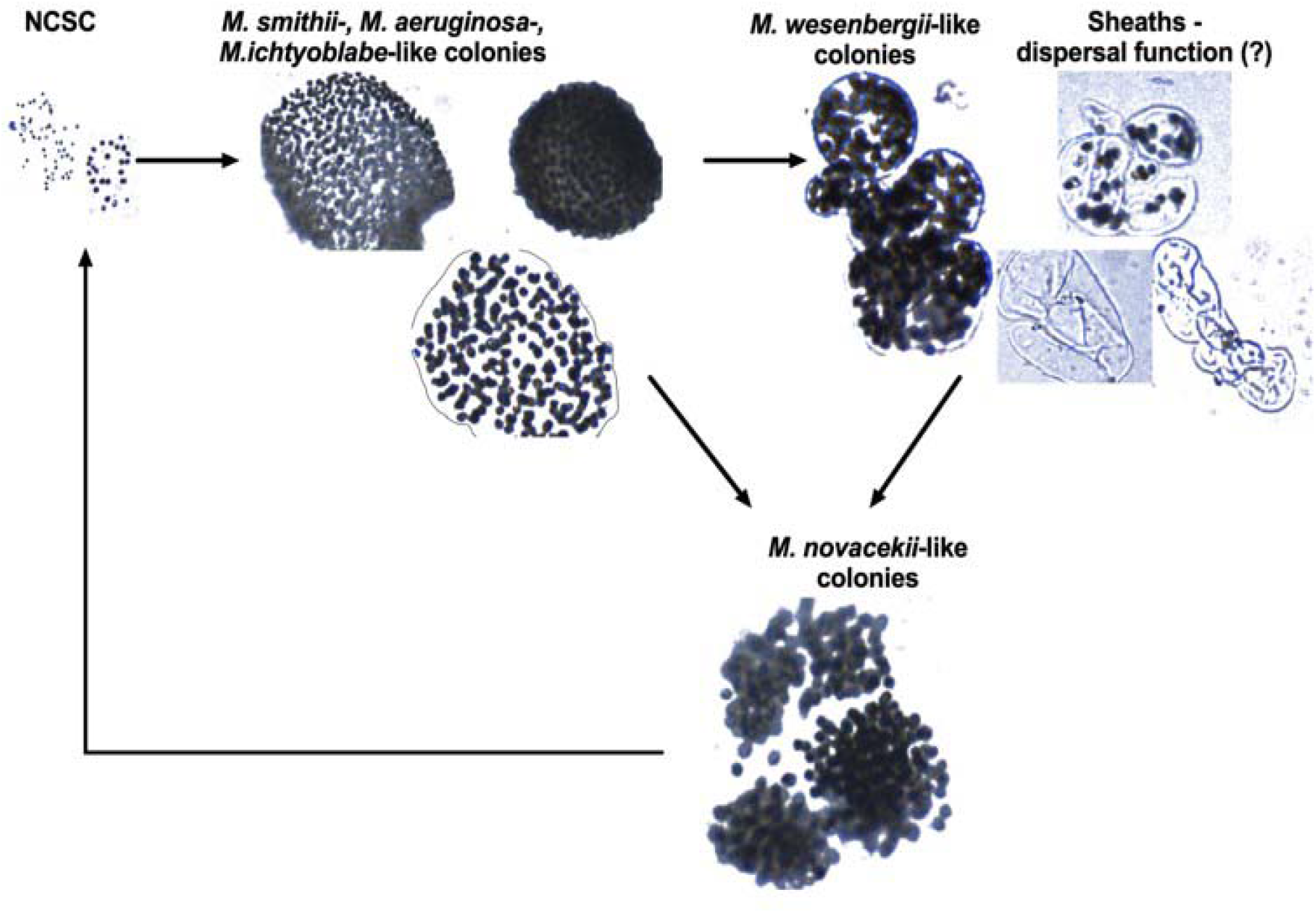
Hypothetical representation of *Microcystis* morphoforms annual cycle with sheaths mirroring *M. wesenbergii* colonial structure and having a role in cyanobacteria dispersal.

Overwintered *Microcystis* cells seem to exist in pre-colonial form and to be the basis for the development of *M. ichtyoblabe, M. smithii,* and *M. aeruginosa* colonies in spring. The next transition during summer is presumably leading to a bloom of *M. wesenbergii* colonies and a bloom of sheaths (some of them still contain *M. wesenbergii* cells, and a structure of sheaths is mirroring *M. wesenbergii* colonies).

### 4.3 Colonial sheaths and their possible role in Microcystis dispersal

The detailed ultrastructure of *Microcystis* colonies was described in 1975 by Kessel and Eloff [58] (in *M. marginata*). The mucilaginous envelope around *M. wesenbergii* colonies, consisting of exopolymeric chains of polysaccharides, decreases their density and might raise the sinking rate [56]. *M. aeruginosa*, *M. ichtyoblabe*, and *M. smithii* also demonstrate mucilage structure enclosing colonies; however, it is less rigid compared to *M. wesenbergii* and sheaths. As Reynolds [5] emphasized, such a secondary structure might be advantageous for numerous reasons. First, mucilage may contribute to buoyancy motility within nutrient- and light-gradient layers of a water body; however, the colony’s size is another prominent factor that dictates the sinking rate. Though the chemical content of *Microcystis* mucilage was studied, the observations of colonial sheaths were reported as a part of *Microcystis* colonies and never as separate entities.

The sheaths are developing by extortion of cells outside or by the death of some *Microcystis* in the colonies. Colonial sheaths appear in significant numbers following *M. wesenberghii* blooming. They may reach significant numbers (thousands) and often still contain few single *Microcystis* cells. We assess such bloom as a specialization for the spreading and dispersal of *Microcystis*. Early Reynolds [5, 56] pointed out the importance of the buoyancy ability of *Microcystis* colonies leading to decreasing colony density. In terms of spreading and expanding in new freshwater aquatic systems, water birds can serve as vectors for dispersal of *Microcystis* colonies with a low density, and colonial sheaths containing *Microcystis* cells means for such dispersion as shown for other aquatic pro- and microeukaryotic communities [57]. Those cells that remain in the original water reservoir can be organized into colonies of *M. novacekii* for overwintering purposes and start the new spring cycle with NCSC. Such results allow us to speculate that *Microcystis* has an enormous advantage due to the coordinative cooperation of its colonial forms on a level higher than a competition of individual colonies more similar to a biofilm-like microbial community.

## 5. Conclusions

To the best of our knowledge, the study is the first to use FlowCAM-based IFC to portray the whole period of *Microcystis* spp. bloom development from colonial non-identified small clusters to colonial sheaths formation and overwintering forms. Our study highlights how IFC analysis can track changes in the intricate development of potentially toxic *Microcystis* spp. population in freshwater reservoirs during the most active phase of their growth. We suggest a hypothetical scheme of transition between *Microcystis* morphoforms. Collectively, the results of this study may add to the knowledge about a framework in which colonial *Microcystis* morphospecies interact with each other on the level of coordinated community and, depending on environmental conditions, thrive in freshwater aquatic systems.

## Author Contributions

A.Z. and Y.M. performed experimental work and wrote an initial draft; D.V.M. performed microscopic taxonomic identification and manuscript editing; T.A.D., E.E.L., E.J., N.S.B.– provided resources, project administration, manuscript editing; N.S.B. – conceptualization. All authors have read and agreed to the published version of the manuscript.

## Funding

This research was funded by MES Kazakhstan, grant number AP14872028 to N.S.B., a Transnational Access granted to N.S.B. through AQUACOSM project (#IFCPHYTO and #SCPCRTNY), by AnaEE Denmark (anaee.dk), the TÜBITAK and program BIDEB2232 (project 118C250) to E.J., and by the European Commission EU H2020-INFRAIA-project (No. 731065) to T.A.D.

## Data Availability Statement

Data available from corresponding author upon reasonable request. The sequencing data for mcyE gene was submitted to DDBJ/ENA/GenBank database and has been deposited under accession numbers OR183428, OR183429, OR183430, OR183431, OR183432, OR183433, OR183434.

## List of abbreviations

AMB: ambient water temperature
ABD: area-based diameter
DNA: deoxyribonucleic acid
Std Dev: standard deviation
EPS: extracellular polymeric substance
FCM: flow cytometry
HABs: harmful algal blooms
IFC: imaging flow cytometry
IPCC: Intergovernmental Panel on Climate Change
LMWE: Lake Mesocosm Warming Experiment
MCs: microcystins
NCSC: small colonial clusters of Microcystis spp.
PCA: Principal Component Analysis
PCR: polymerase chain reaction

## Acknowledgments

We would like to acknowledge help from Veronika Dashkova, Shynar Akhmetova, Aigul Kussanova, Vladimir Novokhatsky, Polina Len and other members of Dr. Barteneva laboratory. Also, we are very grateful to Ann Lene Vigh, Kathrine Tabermann Uhrenholt, and Ivan Nielsen from Aarhus University (Denmark) for assistance with sample collection and measurements of environmental variables. We would like to thank Harry Nelson (ME, USA) for his help and advice.

## Conflicts of Interest

The authors declare no conflict of interest.

## Notes

### Competing Interest Statement

The authors have declared no competing interest.

