## Supplemental_Data_1 for "From colonial clusters to colonial sheaths: analysis of *Microcystis* morphospecies in mesocosm by imaging flow cytometry"

[#]- Current affiliation: IMIM, Graduate School Medical Sciences, HPC FA33 Hanzeplein 1, 9713 GZ, Groningen, The Netherlands

### Corresponding author:

Natasha S. Barteneva;

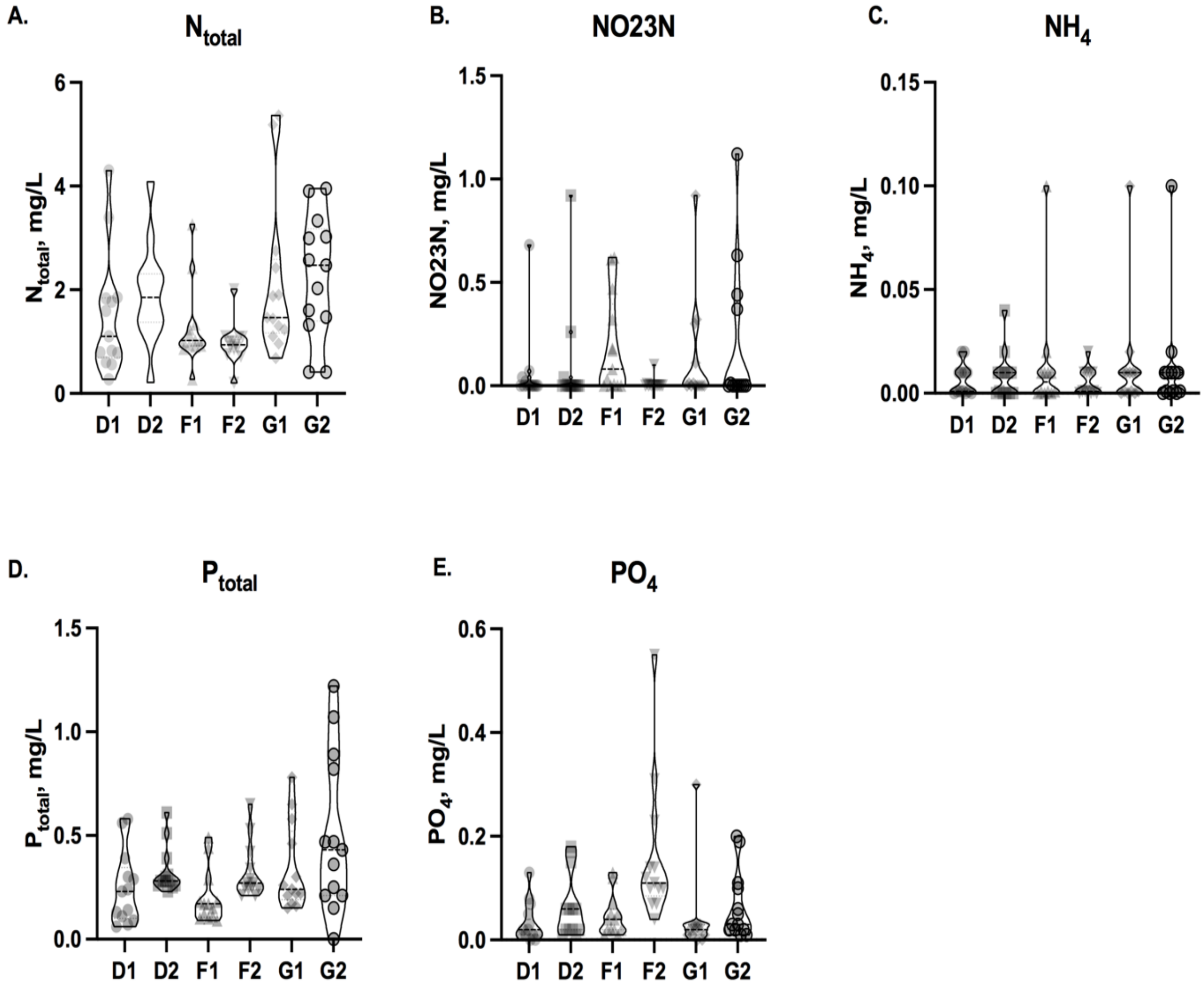


**Supplemental Figure 1**. Concentrations of nutrients in mesocosm tanks. **A**. $N_{total}$; **B.** $NO_{2+3}$ N; **C.** ammonium NH4; **D**. Ptotal; **E**. orthophosphate PO4 in LMWE-2019 mesocosm tanks D1, D2, F1, F2, G1, and G2. Whiskers shows distribution of values from minimum to maximum. Dotted line represents the mean.

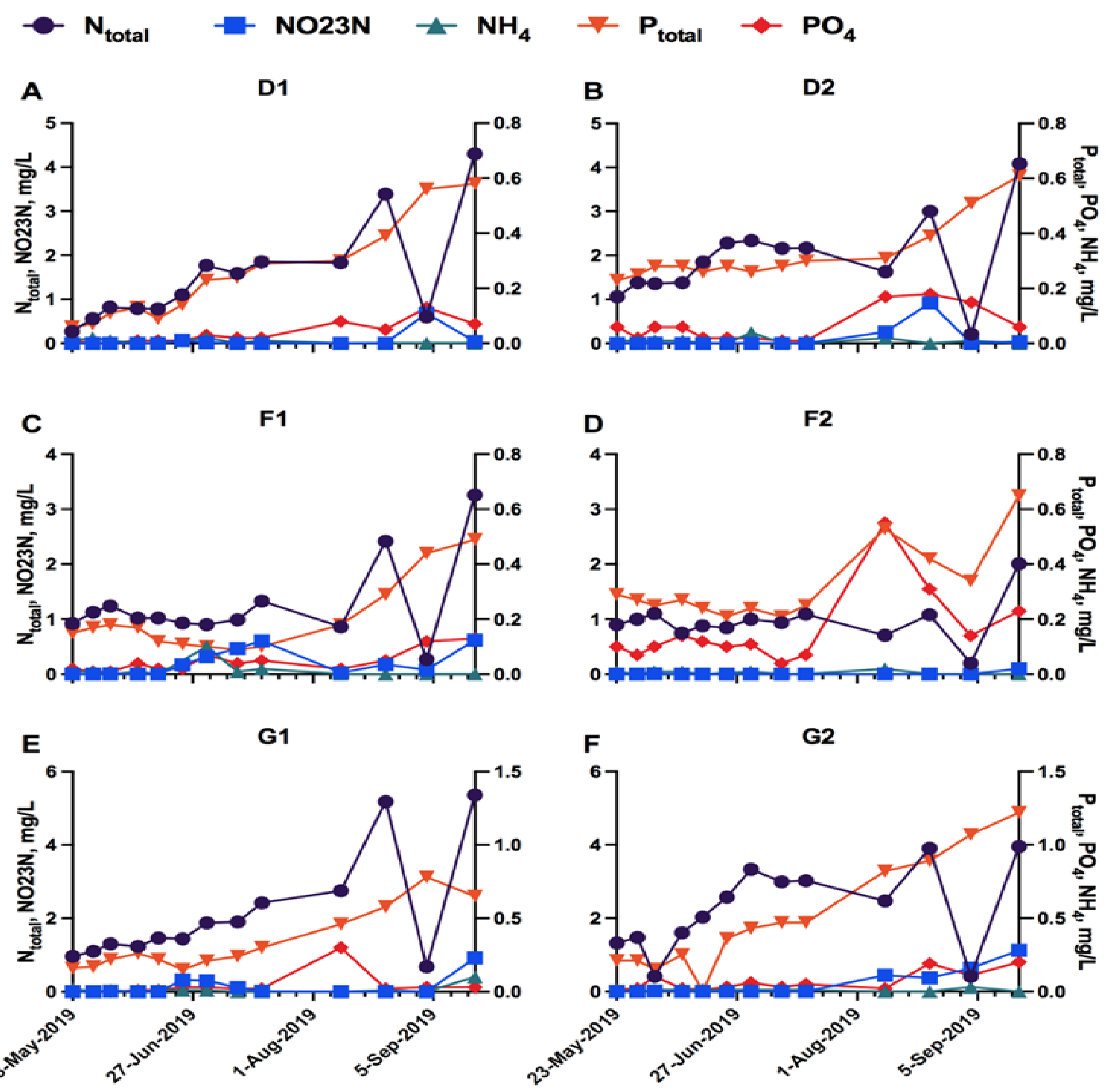


**Supplemental Figure 2**. Temporal variations of nutrients in mesocosm tanks. **A**. tank D1; **B**. tank D2; **C.** tank F1; **D.** tank F2; **E.** tank G1; **F**. tank G2. Measured concentrations of nutrients include total nitrogen ($N_{total}$), nitrate + nitrite (NO2+3N), ammonium (NH4), total phosphorous ($P_{total}$), orthophosphate (PO4). Left Y axis reflects changes in concentration of $N_{total}$ (purple circle) and NO2+3N (blue square), whereas Right Y axis represents seasonal variations of $P_{total}$ (orange inverted triangle), PO4 (red rhombus), and NH4 (teal triangle) concentrations

PCR products were cleaned up and sequenced by Evrogen company. Sequence data is summarized in **Table S1** below. Some samples didn't produce any sequencing data (labeled as "no data").

**Table S1.** The list of DNA sequences for each PCR product from LMWE-2019 mesocosm samples:

| Tank | *Microcystis*-specific mcyE |
|---|---|
| A1 | AAACGATACGCTCATACAGGCTCCAGACYTCATTAGCTTCKGAK<br>GAGAAAAAACCTTTMGAWCCGGCGATKAGGCAGCCACTGAAGG<br>AAGCATTGAGTTTATGGGACGAAAAGATAATCAGCAAGATGTTC<br>ATTACCGTAATAACCTGCCGAGTCTGC |
| D1 | GGTGTGCTCTACCACCCGAATGACTCAGAAATTTAAACCTAGCTT<br>TCTTGATGAGACAAAAACTCTCTTTAGAACCGGCGATTTAGGCAA<br>GCAAACTGCTCCCGGTATCATTGAGTTTATGGGACGAAAAGATA<br>ATCAAGTTAAGGTCAATGGTTATCGAATTGACCCCGGAGAAATTG<br>AATATCAATTGACTCGTTATGCTCCCATTAAAAT |
| F1 | CGACATGCTCTACGACCCGAATGACTCAGAAAATTTAAACCTAGC<br>TTTCTTGATGAGACAAAAACACTCTTTAGAACCGGCGATTTAGGC<br>AAGCAAACTGCTCCCGGTATCATTGAGTTTATGGGACGAAAAGA<br>TAATCAAGTTAAGGTCAATGGTTATCGAATTGACCCCGGAGAAAT<br>TGAATATCAATTGACTCGTTATGCTCCCATTGA |
| G1 | GGAAGCACATACGACCCGAATGACTCAAGAAAAATTTAAACCTA<br>GCTTTCTTGATGAGACAAAAACTCTCTTTAGAACCGGCGATTTAG<br>GCAAGCAAACTGCTCCCGGTATCATTGAGTTTATGGGACGAAAA<br>GATAATCAAGTTAAGGTCAATGGTTATCGAATTGACCCCGGAGA<br>AATTGAATATCAATTGACTCGTTATGCTCCCATTGA |
| A2 | GTGCTCGATGCGCTCTCTCATCCTTGAGTCACTATTAGTTTCTCCA<br>ATGGTATATCAAATCCTTGGATAAAAAGACAGTACAAATTGTATT<br>GCTATTACTTTTAGTGACTGTATTGCAATTTTTTAGAAATCCAAAG<br>AGACCTTTAGAAGTAAACGATTACAATATCATTGCTCCAGTAGAT<br>GGAAAAGTGGTAGTAATTGAAGAAGTTTTTGAACCTGAATATTTT<br>AAAGACCAACGTTTACAAGTTTCTATCTTTATGTCGCCAATAAAT<br>GTACACGTAACTCGTTATGCTCCCATTGA |
| D2 | CAAATCCTCTTGCCACCCGAATGACTCAGAGAATTTAAACCTAGC<br>TTTCTTGATGAGACAAAAACTCTCTTTAGAACCGGCGATTTAGGC<br>AAGCAAACTGCTCCCGGTATCATTGAGTTTATGGGACGAAAAGA<br>TAATCAAGTTAAGGTCAATGGTTATCGAATTGACCCCGGAGAAAT<br>TGAATATCAATTGACTCGTTATGCTCCCATTGATAA |
| F2 | no data |
| G2 | ACGATGATCGCACTAGAGACCATATGATCGAGAAACTTTACACTT<br>AGCTTTCTTAGATGAGACAAAGGCACGTGAGAACCGGCGATTTA<br>GGAAGCAAACTGCTCCCGGAAACATTGAGTTTATGGGACGAAAA<br>GATAATCAAGTTAAGGTCAATGGTTATCGAATTGACCCCGGAAAT<br>TGAAAAATTGTTCGTTGTCCTTGA |

Then these sequences were searched by NCBI blastn program and the most similar matches are listed below in **Table S2.** The sequencing results shows presence of *Microcystis*-specific mcyE gene in samples from all mesocosm tanks, except for A2 and F2 tanks.

**Table S2.** Alignment matches of mesocosm samples with NCBI Nucleotide collection database:

| Sample-target names | Size (bp) | Description of match | Max Score | E value | Percent ident |
|---|---|---|---|---|---|
| A1-mcyE | 158 | Uncultured *Microcystis* sp. microcystin synthetase E (mcyE) gene, partial cds | 58.4 | 4e-04 | 100% |
| D1-mcyE | 213 | *Microcystis aeruginosa* FCY-26 MсyE (mcyE) gene, complete cds | 344 | 4e-90 | 97.10% |
| F1-mcyE | 212 | *Microcystis aeruginosa* FCY-26 MсyE (mcyE) gene, complete cds | 346 | 1e-90 | 98.01% |
| G1-mcyE | 213 | *Microcystis aeruginosa* FCY-26 MсyE (mcyE) gene, partial cds | 364 | 3e-96 | 99.50% |
| A2-mcyE | 301 | *Flavobacterium haoranii* strain KCTC 23008 chromosome, complete genome | 159 | 2e-34 | 81.96% |
| D2-mcyE | 215 | *Microcystis aeruginosa* FCY-26 MсyE (mcyE) gene, complete cds | 346 | 1e-90 | 98.01% |
| F2-mcyE | no data | | | | |
| G2-mcyE | 202 | *Microcystis aeruginosa* FCY-26 MсyE (mcyE) gene, complete cds | 195 | 4e-45 | 88.55% |
